# U-RISC: an ultra-high-resolution electron microscopy dataset challenging existing deep learning algorithms

**DOI:** 10.1101/2021.05.30.446334

**Authors:** Ruohua Shi, Wenyao Wang, Zhixuan Li, Liuyuan He, Kaiwen Sheng, Lei Ma, Kai Du, Tingting Jiang, Tiejun Huang

## Abstract

Connectomics is a developing field aiming at reconstructing the connection of the neural system at nanometer scale. Computer vision technology, especially deep learning methods used in image processing, has promoted connectomic data analysis to a new era. However, the performance of the state-of-the-art methods still falls behind the demand of scientific research. Inspired by the success of ImageNet, we present the U-RISC, an annotated **U**ltra-high **R**esolution **I**mage **S**egmentation dataset for **C**ell membrane, which is the largest cell membrane annotated Electron Microscopy (EM) dataset with a resolution of 2.18nm/pixel. Multiple iterative annotations ensured the quality of the dataset. Through an open competition, we reveal that the performance of current deep learning methods still has a considerable gap with human-level, different from ISBI 2012, on which the performance of deep learning is close to human. To explore the causes of this discrepancy, we analyze the neural networks with a visualization method, attribution analysis. We find that in U-RISC, it requires a larger area around a pixel to predict whether the pixel belongs to the cell membrane or not. Finally, we integrate currently available methods to provide a new benchmark (0.67, 10% higher than the leader of competition, 0.61) for cell membrane segmentation on U-RISC and propose some suggestions in developing deep learning algorithms. The U-RISC dataset and the deep learning codes used in this paper will be publicly available.

## Introduction

Accurate descriptions of neurons and their connections are fundamental to modern neuroscience. By depicting neurons with the help of Golgi-staining method (***Golgi, 1885***)), Cajal could propose the classic “Neuron Doctrine” more than a century ago (***y Cajal, 1888***), which opened a new era for modern neuroscience. Nowadays, the development of electron microscopy (EM) has enabled us to further explore the structural details of the neural system at nanometer (nm) scales (***Kornfeld and Denk, 2018; Shawn, 2016***) opening up a new field called “Connectomics” that aims to reconstruct every single connection in the neural system. One milestone of connectomics is the *C. elegans* project (***White et al., 1986***) which maps all 302 neurons and 7,000 connections in a worm. Recently, a small piece of human cortex was imaged with a high-speed scanning EM, which maps ~50,000 neurons and ~110,000,000 synaptic connections (***Shapson-Coe et al., 2021***). Connectomic data increases exponentially with a higher resolution of EM and a larger neural tissue volume, even reaching petabyte (PB) scale (***Shapson-Coe et al., 2021***). Just as it took almost 15 years to complete the connectome of *C. elegans*, structural reconstruction for higher-level creatures is becoming more and more daunting with the explosion of connectomic data. Among many bottlenecks, accurate annotation from large amounts of EM images is the first one that has to be solved.

Manual annotation of all the connectomic data is infeasible because of the high annotation cost. To reduce the burden of manual annotation for humans, one would hope to enable a machine to annotate the connectomic data with near-human performance automatically. Hopes are higher today because of the rapid development of deep learning methods. However, even with deep learning, it still requires tremendous efforts to achieve human-level performance on this challenging task. There were a few successful experiences to learn from the computer science community to make the deep learning method fully comparable to humans in connectomics. The success of deep learning methods highly depends on the amount of training data and the quality of annotation. Take the task of image classification as an example; ImageNet (***Russakovsky et al., 2015***) has set up a research paradigm of applying deep learning methods for vision tasks. In 2009, by releasing a large-scale accurately annotated dataset, ImageNet provided a benchmark (72%) for image classification. From 2010 to 2017, a challenge called “The ImageNet Large Scale Visual Recognition Challenge (ILSVRC)” was organized every year. This challenge significantly boosted the development of deep learning algorithms. Many champions of this challenge have become the milestones of deep learning methods, such as AlexNet (***Krizhevsky et al., 2012***), VGG (***Simonyan and Zisserman, 2014***), GoogleNet (***Szegedy et al., 2015***), ResNet (***He et al., 2016***), etc. As shown in ***Figure 1***, deep learning performance on image classification finally exceeded human level (95%) after eight years of development. To summarize, there is a roadmap for the success of ImageNet, which includes three key steps: the first step is to establish a large-scale dataset with high-quality annotation, which is very important for deep learning. Based on the dataset, the second step is organizing a challenge that can evaluate algorithms at a large scale and allow researchers to estimate the progress of their algorithms, taking advantage of the expensive annotation effort. The third step is the design of new algorithms based on the previous two steps. Each of the three stages is indispensable.

**Figure 1.**
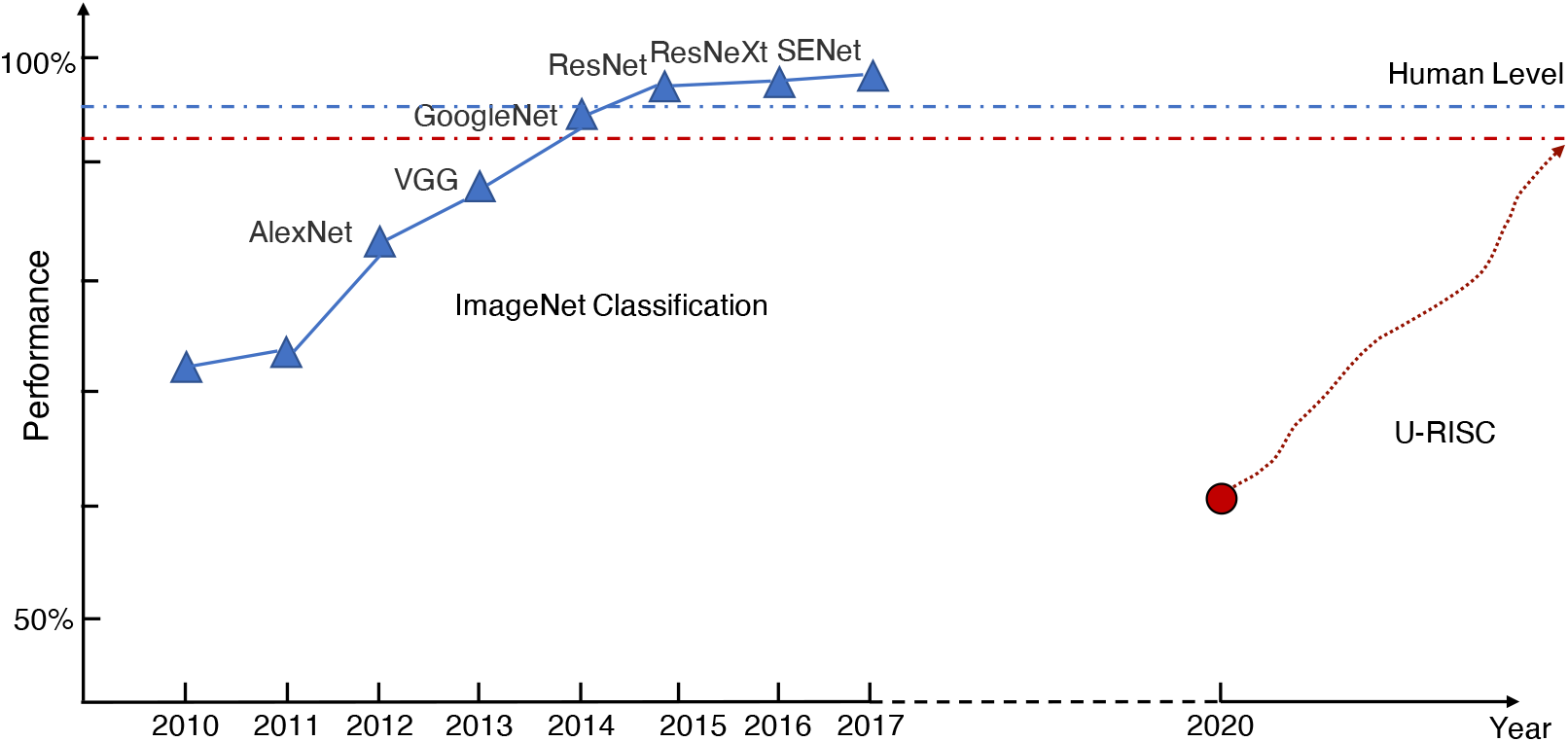
The history of ImageNet.

Following the success of ImageNet, significant progress of EM automatic segmentation was achieved by the 2012 IEEE International Symposium on Biomedical Imaging (ISBI 2012), which was the first challenge on EM automatic segmentation with releasing a publicly available dataset (***Arganda-Carreras et al., 2015***). The state-of-the-art (SOTA) method exhibited unprecedented accuracy in EM cellular segmentation on the dataset of ISBI 2012. In particular, the deep learning method “U-Net” (***Ronneberger et al., 2015***), which was first proposed during the challenge, becomes the backbone of many SOTA methods in the field. However, today many deep learning methods have become “exceedingly accurate” and likely saturated at the ISBI 2012 (***Arganda-Carreras et al., 2015***). In addition, ISBI 2012 images are 512 × 512 pixels with a resolution of 4 nm/pixel × 4 nm/pixel, while there are many EM images with higher resolution in connectomics because enough high resolution is essential to unravel the neural structures unambiguously. For instance, 2nm has been suggested as the historical “gold standard” to identify synapses (***DeBello et al., 2014***), in particular to identify gap junctions (***Leitch, 1992***) which are common in neural tissues (***Anderson et al., 2009***). It is not clear if previous classic deep learning methods developed on EM images with relatively lower resolution can still work well on datasets with higher resolutions.

Here, to promote the deep learning algorithms in EM datasets, we initiate a new roadmap: We first annotated the retinal connectomic data, RC1, from rabbit(***Anderson et al., 2011***) and presented a brand new annotated EM dataset named U-RISC (Ultra-high Resolution Image Segmentation dataset for Cell membrane). Compared with ISBI 2012, U-RISC has a higher resolution of 2.18 nm/pixel and a larger size of 9958 × 9959 pixels. The precision of annotation was ensured by multi-steps of iterative verification, costing over 10,000 labour hours in total. Next, based on U-RISC, a competition of cellular membrane prediction was also organized. Surprisingly, from 448 domestic participants/teams, the top performance of deep learning methods on U-RISC (~ 0.6, F1-score) was far below the human-level accuracy (> 0.9), in contrast with the near-human performance of deep learning methods in ISBI 2012. We then made fair comparisons between ISBI 2012 and U-RISC with the same segmentation methods, including U-Net. The comparison results confirmed that U-RISC indeed provides new challenges to existing deep learning methods. U-Net, for example, dropped from 0.97 in ISBI 2012 to 0.57 in U-RISC. To further explore how these methods work on segmentation tasks, we introduced a gradient-based attribution method, integrated gradient (***Sundararajan et al., 2017***), to analyze ISBI 2012 and U-RISC. The result showed that when deciding on whether a pixel belonged to the cell membrane or not, deep learning methods represented by U-Net would refer to a larger attribution region on U-RISC (about four times on average) than that on ISBI 2012. It suggests that the deep learning methods might require more background information to decide the segmentation of the U-RISC dataset. Finally, we integrated current available advanced methods, combining U-Net and transfer learning recently introduced (***Conrad and Narayan, 2021***), and provided a benchmark (0.6659), about 10% higher than the leader board (0.6070), for U-RISC.

Overall, our contribution in this work lies mainly in the following three parts: (1) we provided the community a brand new publicly available annotated mammalian EM dataset with known highest resolution (~ 2.18 nm/pixel) and largest image size (9958 pixel × 9959 pixel); (2) we organized a competition and made a comprehensive analysis to reveal the challenges of U-RISC for deep learning methods; (3) we improved the benchmark with 10% to the F1-score of 0.6659. In discussion, we proposed further suggestions for improving segmentation methods from perspectives of model design, loss function design, data processing, etc. We hope our dataset and analysis can help researchers gain insights into designing more robust methods, which can finally accelerate the speed of untangling brain connectivity.

## Results

### The largest ultra-high-resolution EM cell membrane segmentation dataset

Along with this paper, we proposed a new EM dataset with cell membrane annotated: Ultra-high Resolution Images Segmentation dataset for Cell membrane (**U-RISC**). To our best knowledge, U-RISC has the highest resolution among the publicly available annotated EM datasets (see ***Figure 2***(A) as an example). It was annotated upon the rabbit retinal connectomic dataset RC1 (***Anderson et al., 2011***) with a 2.18 nm/pixel resolution at both *x* and *y* axes. The size of individual image and the total number of images in U-RISC both exceed the published datasets by far, taking ISBI 2012 as an example (***Figure2***(C)) (120 pairs of 9958 pixel × 9959 pixel images in U-RISC and 30 pairs of 512 pixel × 512 pixel images in ISBI 2012). One characteristic of U-RISC is that cell membranes only cover a small area of the images, making it an imbalanced dataset for deep learning (an average of 5.10% ± 2% in U-RISC compared with 21.65%± 2% in ISBI 2012).

**Figure 2.**
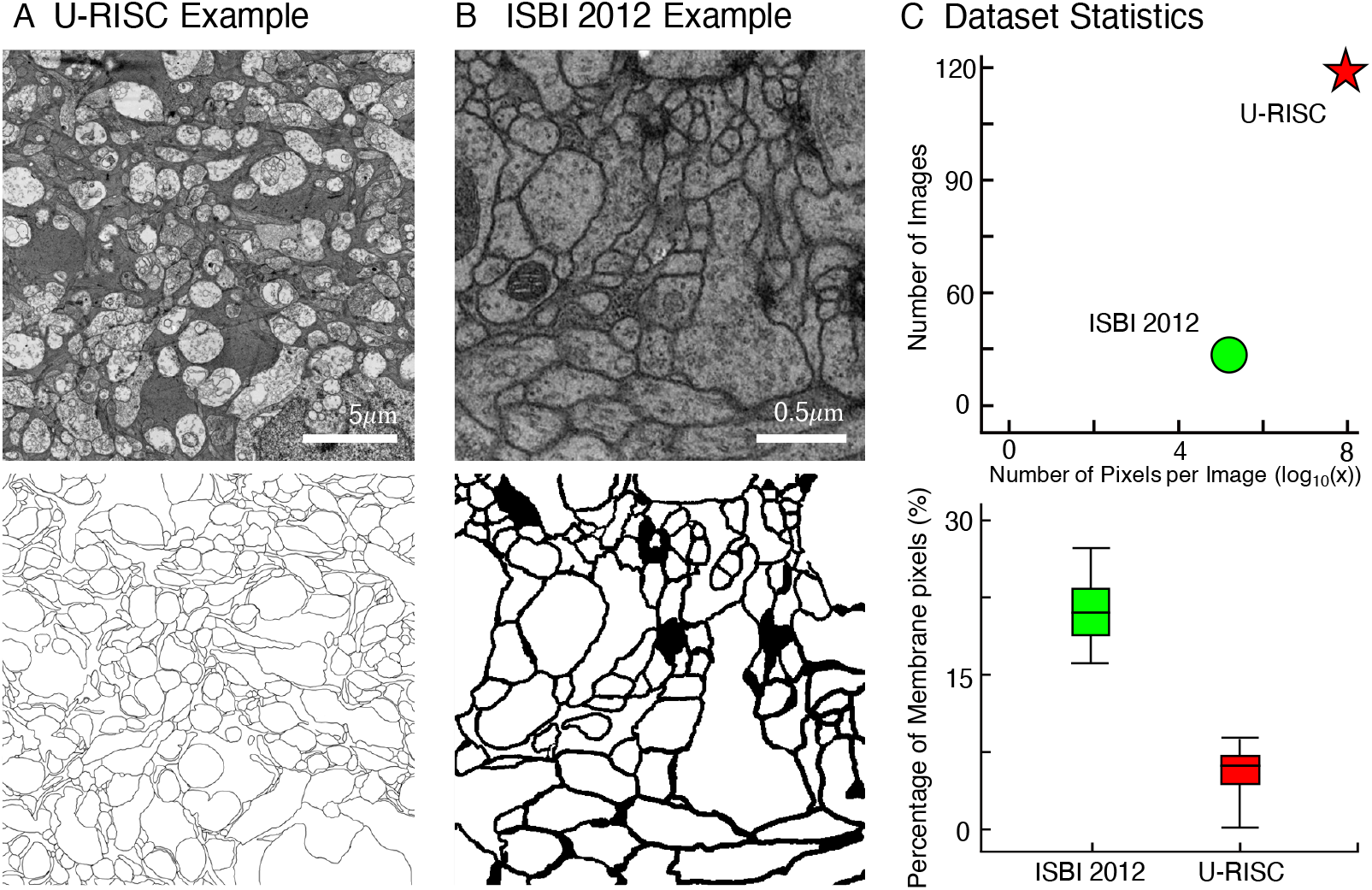
Comparison between U-RISC and ISBI 2012. (A and B) An example of U-RISC and ISBI 2012 data includes the raw EM image (top) and the corresponding annotation result (bottom). Black pixels in annotation results represent cellular membranes. (C) (Top) Both the number and size of images in U-RISC surpass those in ISBI 2012. (Bottom) The proportion of annotated pixels, 5.10%±2% in ISBI 2012 and 21.65%±2% in U-RISC, making latter a more imbalanced dataset.

We employed an iterative manual annotation procedure to ensure the quality of annotation. Because of the difficulty of distinguishing cell membrane from organelle membrane, special attention was paid to exclude organelle membrane from annotation (***Figure 3***(A)). In practical connectomic research, the image quality can be affected by many reasons like insufficient staining, thick section, etc. Considering this, we retained several images with low quality in U-RISC to make the dataset closer to the actual situation. Annotation on these images costs more time and caution (***Figure 3***(B)). Labeling errors could be detected and then corrected in each round of iteration (***Figure 4***). For scientific research reasons, the human labeling process is very valuable for uncovering the human learning process. Therefore, the intermediate annotated results were also reserved for public release (https://brain.baai.ac.cn/biodb-rabbit-details.html).

**Figure 3.**
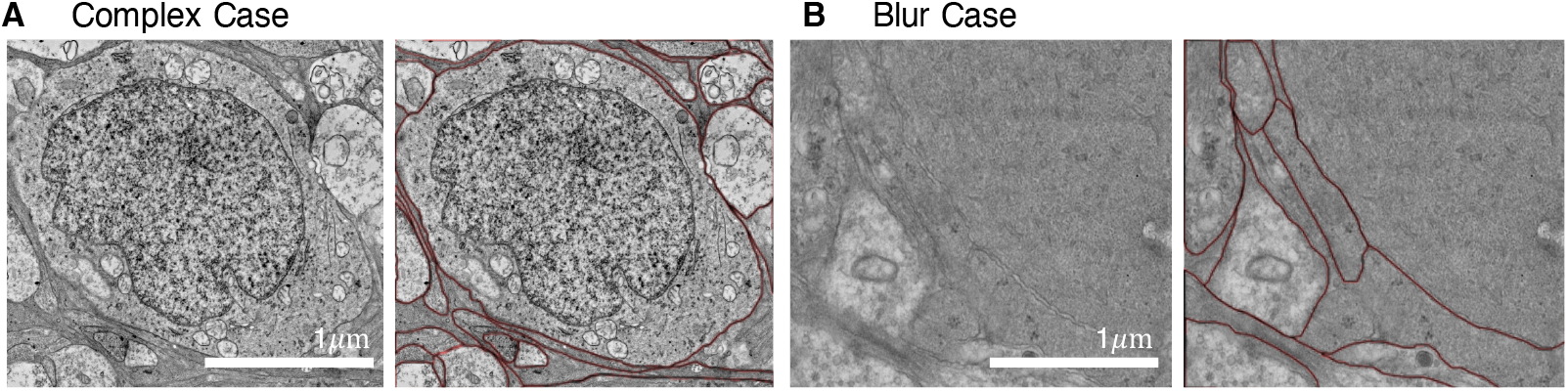
Examples of images with their annotations. (A) Organelle membranes were cautiously avoided to be annotated. (B) More time and patience were needed to annotate the image with low contrast.

**Figure 4.**
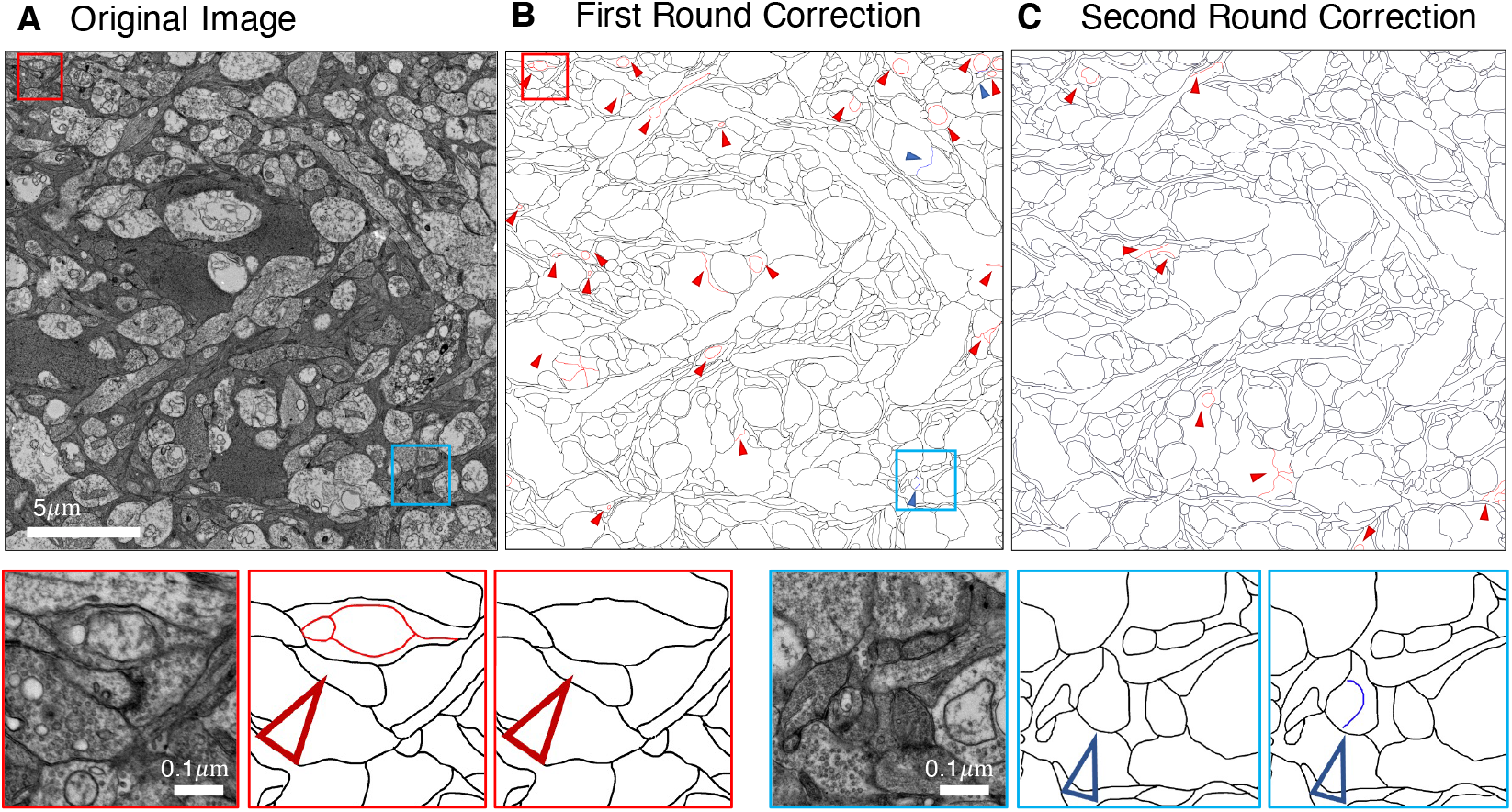
Example of iterative human annotation. (A) Original image to be annotated. (B) Many errors were found out in the first round of annotation. (C) After correction, much fewer errors were detected in the second round of annotation, and the correction results were served as the final annotation. Red small triangles and boxes indicate false positive errors (enlargement in the bottom left), blue for false-negative errors (enlargement in the bottom right).

### Ultra-high resolution EM images segmentation competition

To investigate the performance of the deep learning methods on U-RISC and propose a benchmark, a competition about cellular membrane segmentation was organized by BAAI (Beijing Academy of Artificial Intelligence Institution, Beijing, China) and PKU (Peking University, Beijing, China)^1^. In total, 448 participants took part in the competition, mainly from domestic competitive universities, research organizations, and top IT institutions.

There were two tracks in the competition (***Table 1***): Track 1 used original images with the size of 9958 pixel × 9959 pixel as training and testing datasets. In Track 2, images were downsampled to the size of 1024 pixel × 1024 pixel. The purpose of Track 2 was to allow researchers with limited computational resources to participate in the competition. The final round of human annotation was used as the ground truth to evaluate the algorithms, and an F1-score was applied as the evaluation metric (for details, please see Methods and Materials).

**Table 1.**
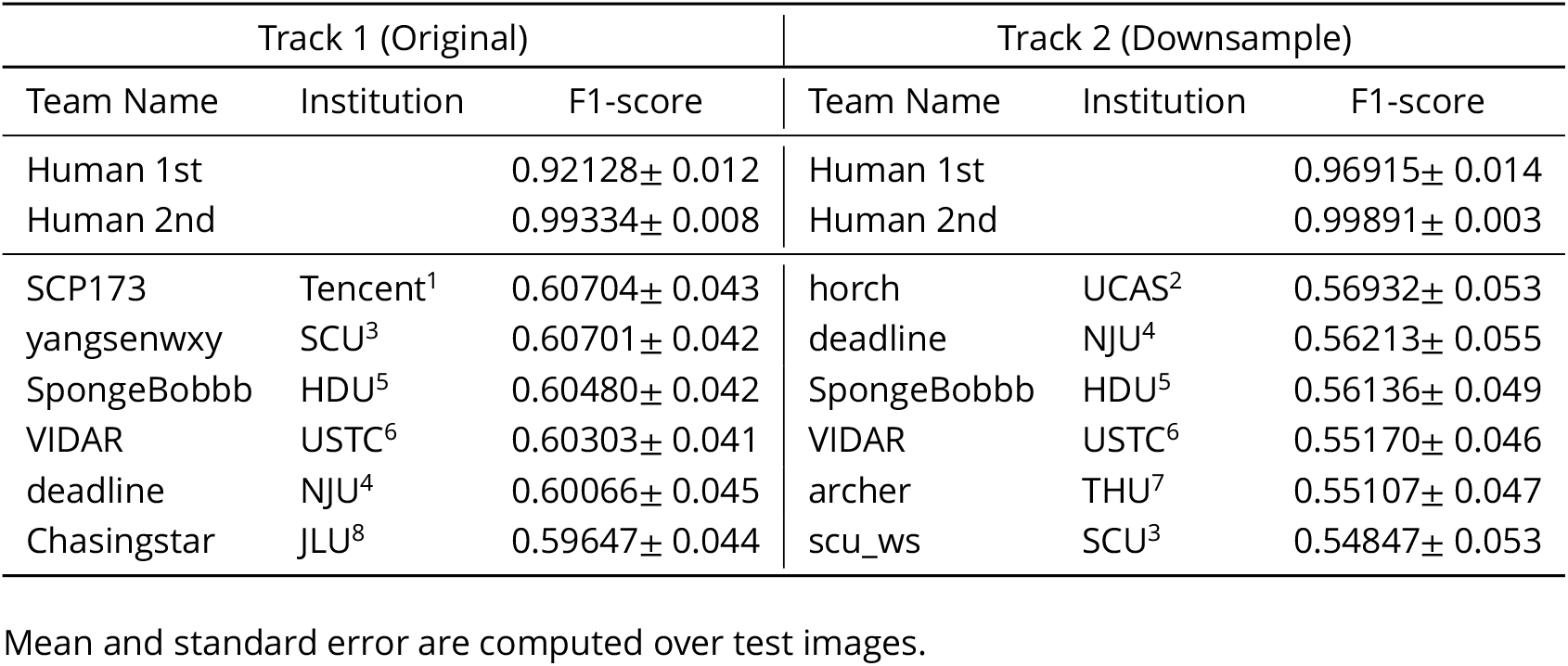
Leaderboard of Track 1 and Track 2.

Surprisingly, from the competition, top 6 teams in each track gained F1-scores around 0.6 on U-RISC, which were far below the human levels (0.92 and 0.99, the first and second rounds of annotation). However, previous research has shown that the performance of the top teams in ISBI 2012 had already been reasonably close to the human level (***Arganda-Carreras et al., 2015***). To investigate causes of the performance gap between the methods and humans on U-RISC, we first surveyed the top 6 teams in our competition. It indicated that a variety of current popular approaches to segmentation were utilized (***Figure 5***). From the choice of models (***Figure 5***(A)), the participants used current popular image segmentation networks, such as U-Net (***Ronneberger et al., 2015***), Efficientnet (***Tan and Le, 2019***) and CASENet (***Yu et al., 2017***). For backbone selection, ResNet (***He et al., 2016***) and their variants were the most chosen architectures. Data augmentation was ubiquitously applied to improve the generalization of the models. About 13% of the participants used Hypercolumns (***Hariharan et al., 2015***) to improve the expressiveness of the model. From the design of loss function, functions that can adjust penalty ratios according to sample distributions were applied to reduce the effect of sample imbalance, such as Dice loss (***Dice, 1945***), Focal loss (***Lin et al., 2017***), and BCE loss (***Cui et al., 2019***). And Adam (***Kingma and Ba, 2014***) was shown to be the most chosen optimization method.

**Figure 5.**
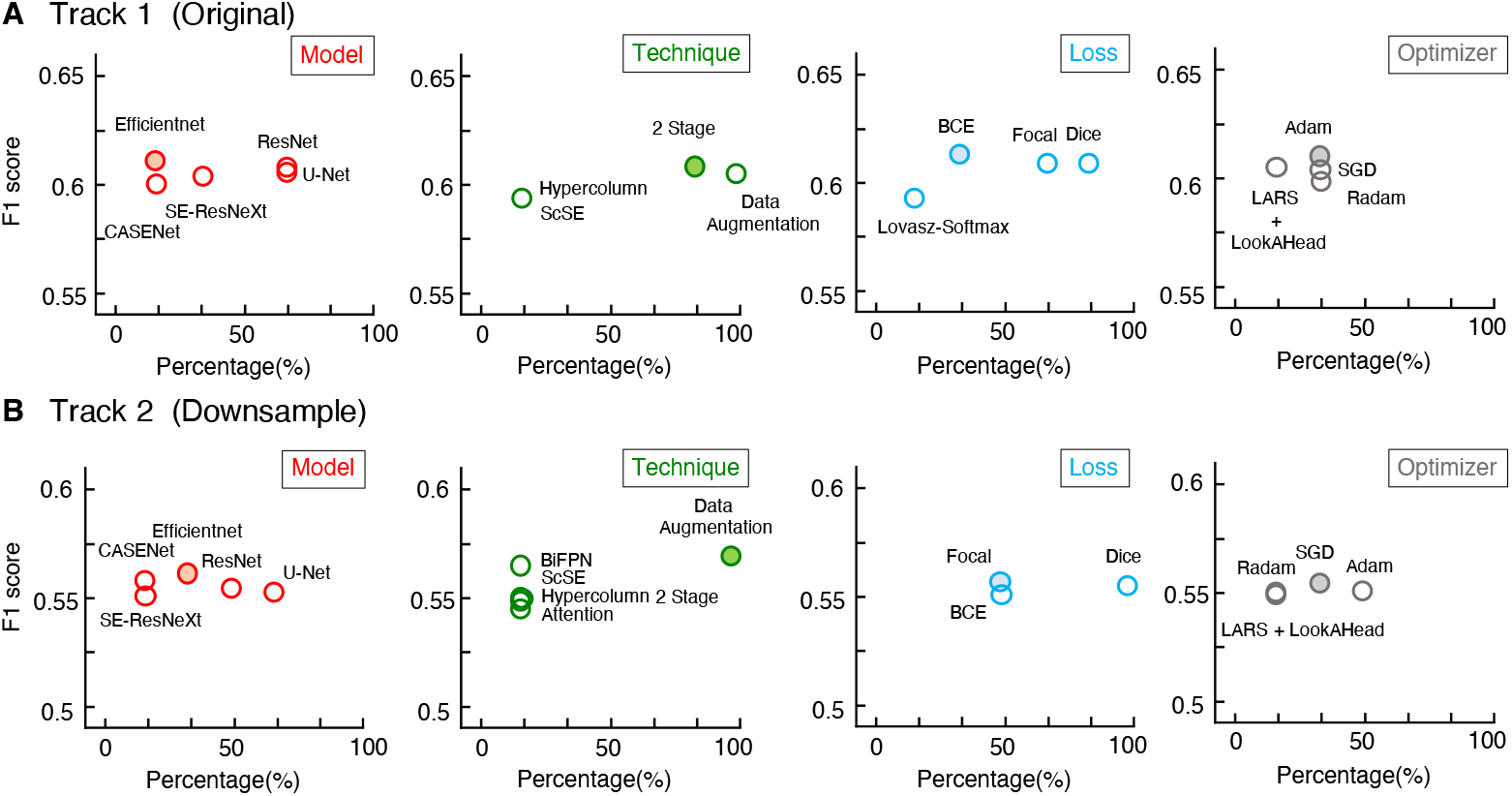
Mean F1 scores of teams with different methods used. (A and B) The statistics of Track 1 and Track 2, respectively. The X-axis represents the proportion of the team with the method, Y-axis represents the average of F1 scores.

The analysis suggested that even though participants had considered many popular methods, their performance was still not satisfactory and varied only slightly between each other. To identify whether this was because of the challenges of U-RISC or the methods themselves, we picked out three widely used methods: U-Net (***Ronneberger et al., 2015***), LinkNet (***Chaurasia and Culurciello, 2017***), and CASENet (***Yu et al., 2017***). We conducted a fair comparison between the performance of each method on U-RISC (Track 1) and ISBI 2012. Results showed that these methods could reach over 0.97 (F1-score) in ISBI 2012, but only between 0.57-0.61 in U-RISC (***Table 2***), which confirmed that the performance gap in competition comes from the challenges of U-RISC.

**Table 2.**
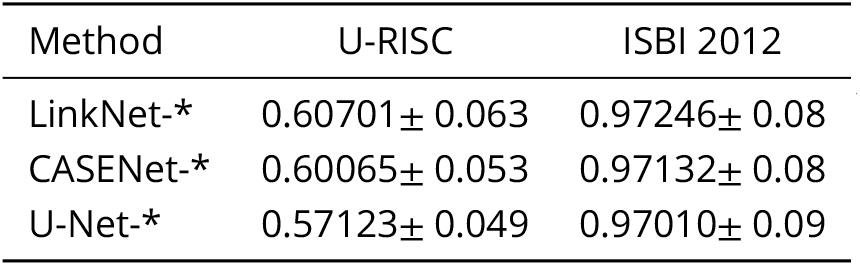
F1-scores in U-RISC and ISBI 2012.

What are the unique challenges brought by U-RISC to deep learning algorithms? Two types. of errors were analyzed first: false-positive errors, which led to incorrect membrane predictions, and false-negative errors, which caused uncontinuity of cell membrane. According to our analysis, both false-positive errors (pink boxes) and false-negative errors (orange boxes) were common in U-RISC, which were rare in ISBI 2012 (***Figure 6***(B) (C)). Further investigations for the networks are required to explore the reason and find ways to reduce the errors.

**Figure 6.**
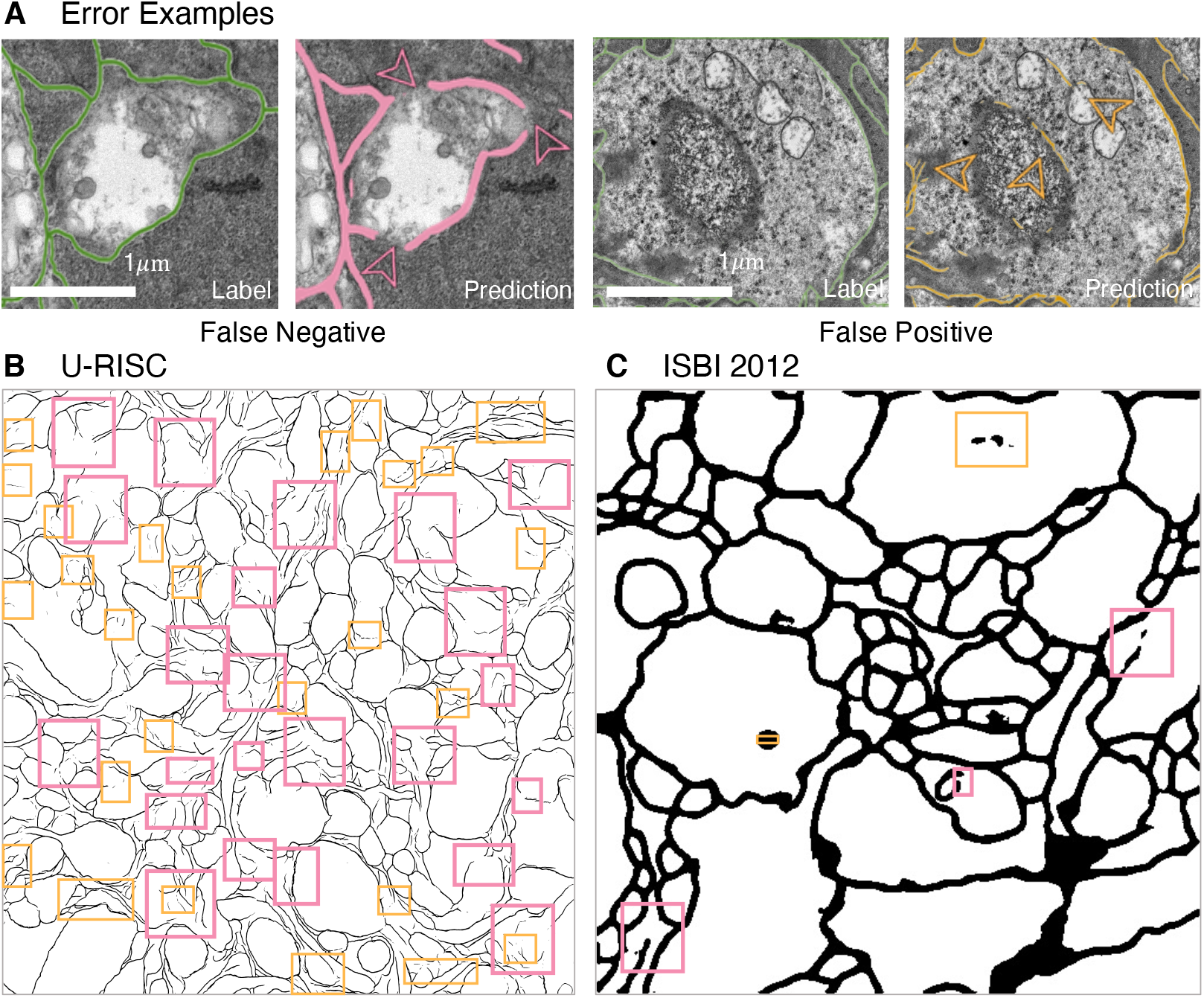
Errors in segmentation predictions of U-RISC and ISBI 2012. (A) The examples of False Positive and False Negative errors. (B and C) The examples of two errors in the segmentations of U-RISC and ISBI 2012. Pink arrows and lines represent false-negative errors, and orange represents false positive errors. **Figure 6-Figure supplement 1.** Supplements for segmentation predictions of U-RISC. **Figure 6-Figure supplement 2.** Supplements for segmentation predictions of ISBI 2012.

### Attribution analysis of the deep learning method on U-RISC and ISBI 2012

To acquire a deeper understanding of the different performances in U-RISC and ISBI 2012, we per-formed an attribution analysis (***Ancona et al., 2019***) on the trained U-Net. We selected the gradient-based attribution method, **integral gradient (IG)** (***Sundararajan et al., 2017***), which is widely applied on explainable artificial intelligence, such as understanding feature importance (***Adadi and Berrada, 2018***), identifying data skew (***Clark et al., 2019***), and debugging model performance (***Guidotti et al., 2018***). In brief, IG aims to explain the relationship between predictions and input features based on gradients (***Figure 7***(A)). The IG output is plotted in **Attribution Fields** to reflect their contribution to the final prediction. In the heatmap, each pixel was assigned with a normalized value between [-1,1]. With IG,we analyzed the attribution field of each predicted pixel of U-Net in U-RISC and ISBI 2012. Color and shade were used to represent the normalized contribution values in attribution fields ***Figure 7***(B). For a fair comparison between U-RISC and ISBI 2012, areas of **Pixel Attribution Fields** *S_k_* were converted to physical size according to their respective resolutions.

**Figure 7.**
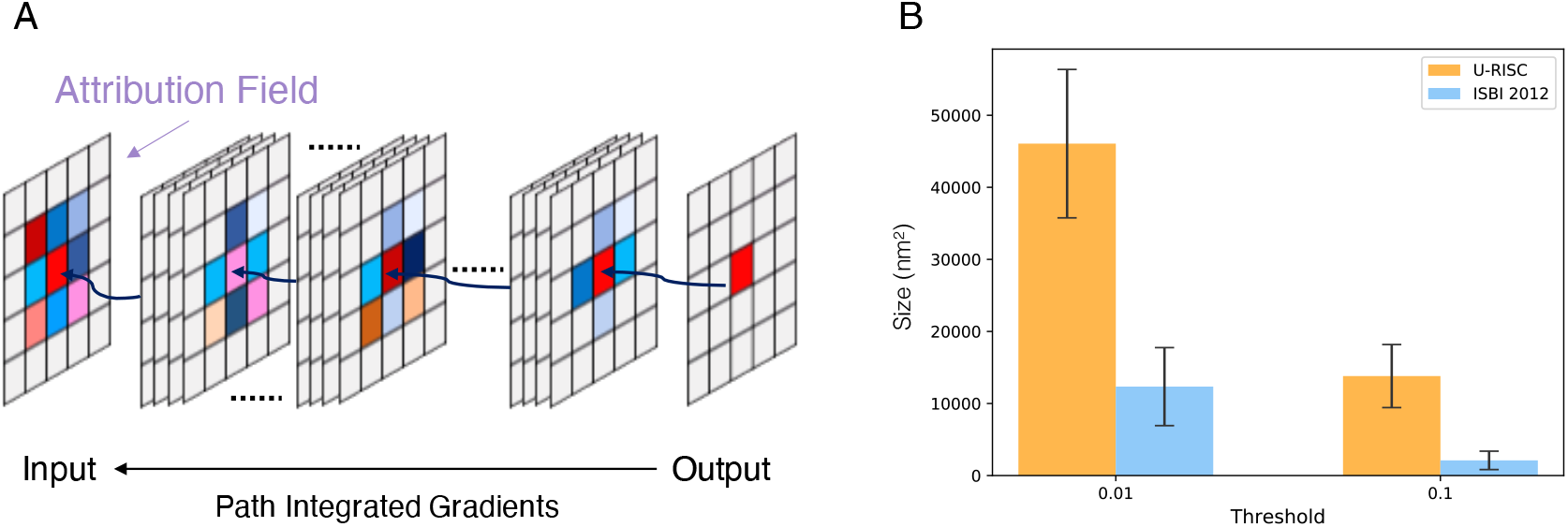
Attribution analysis. (A) Integrated gradients attribution method. (B) Statistics of attribution filed for U-RISC and ISBI 2012.

***Figure 8*** shows the examples of attribution fields, where bounding boxes with different colors represented different pixel classifications, green for a correct predicted pixel, orange for a false positive error, and pink for a false negative error. We noticed that the areas of attribution fields *S_k_* of two datasets were both relatively minor to the whole images (***Figure 7***(B)). For example, at the threshold of *k* > 0.01, the *S_k_* of the correct cases accounted for only 5.1% and 0.8% relative to the whole image (the green bounding boxes in ***Figure 8***). This suggested that U-Net would focus on local characteristics within small areas of the images when making predictions. In addition, we found that the averaged *S_k_* of each predicted pixel in U-RISC was significantly larger than that in ISBI 2012, specifically 46000 *nm*^2^ in U-RISC and 10300 *nm*^2^ in ISBI 2012. Taken together, U-Net would predict cell mambrane according to local information around the pixel, and the average attribution field was larger in U-RISC than that of ISBI 2012. All of these indicate that more information is required for the segmentation in U-RISC.

**Figure 8.**
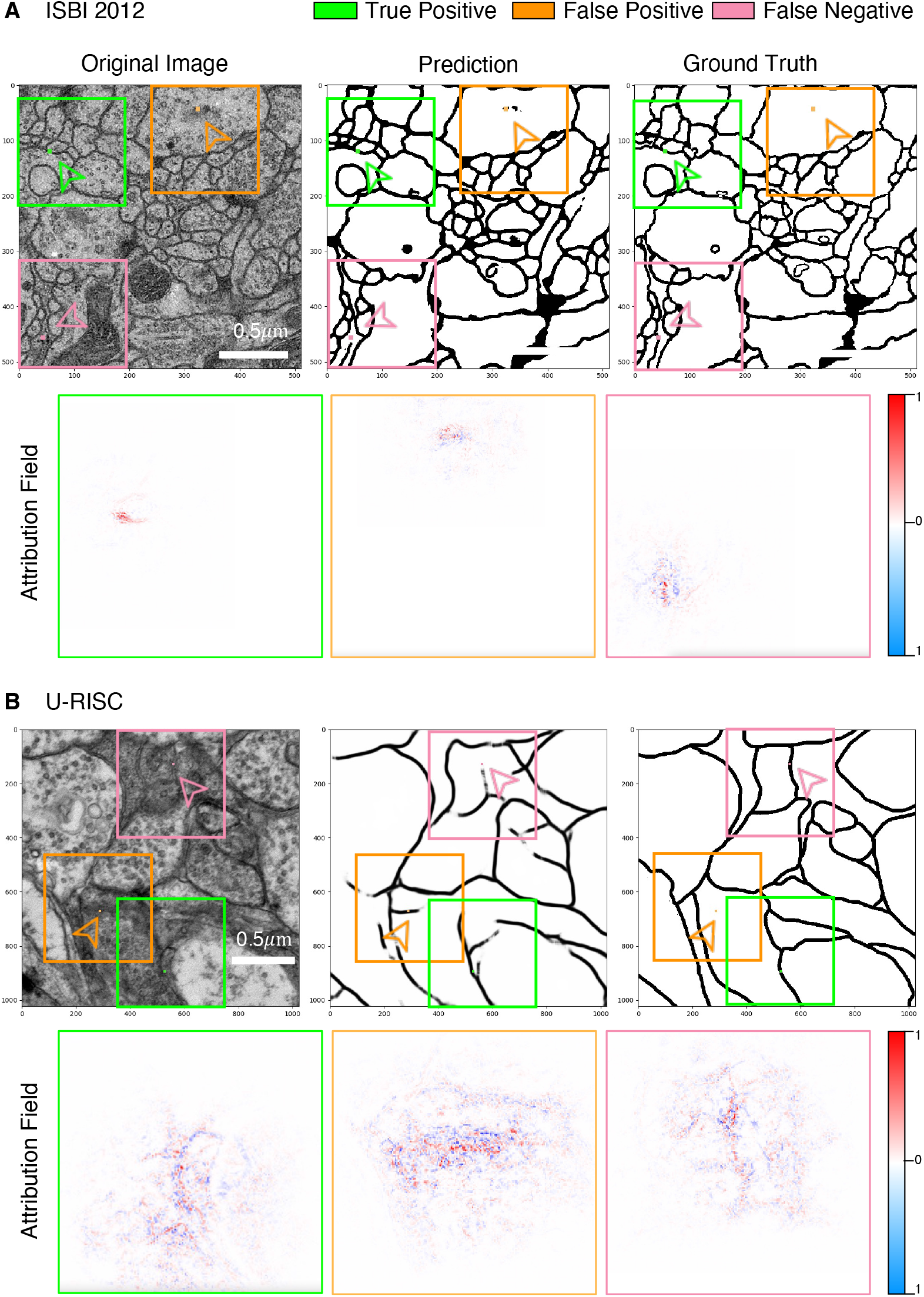
Attribution analysis. (A and B) Attribution fields of ISBI 2012 and U-RISC dataset. The first line represents the original image, network prediction result and annotation respectively. The pixels pointed by green (correct cell membrane pixel), orange (false positive predicted pixel) and pink (false negative predicted pixel) arrows are the prediction points used in attribution method. Images in the three color boxes with the same size in the second line represent the attribution field corresponding to the above three pixels. Blue means the network is likely to predict the pixels as cell membrane, while opposite for red. **Figure 8-Figure supplement 1.** Supplements for attribution analysis on ISBI 2012. **Figure 8-Figure supplement 2.** Supplements for attribution analysis on U-RISC.

### U-Net-transfer model achieve the state-of-the-art result on U-RISC benchmark

Considering both the comprehensive analyses of competition and attribution analysis, we integrated outperformed methods to develop our method (***Figure 9***(A)). For basic segmentation architecture, we chose U-Net due to its better characteristic extraction ability. Many valuable techniques were also considered, including cross-crop strategy for saving computational resources and data augmentation to increase data diversity. We chose both focal loss and dice loss to deal with the imbalance of samples for loss function design. Some parameters used for training were also optimized, such as batch-size/GPU (4) and the number of GPUs (8). For more details, please refer to the part of Segmentation Networks in Methods and Materials. Especially, recent research has shown that transfer learning with domain-specific annotated datasets could be effective in elevating deep learning models’ performance (***Conrad and Narayan, 2021***). Therefore, we introduced a pre-trained model, trained with MoCoV2 (***He et al., 2020***) on CEM500K (***Conrad and Narayan, 2021***). The segmentation result showed that the F1 scores of our method were 10% higher than the leader of competition (0.66 vs 0.61 in ***Table 2***). Thus we provide a new benchmark on the cellular membrane segmentation of U-RISC.

**Figure 9.**
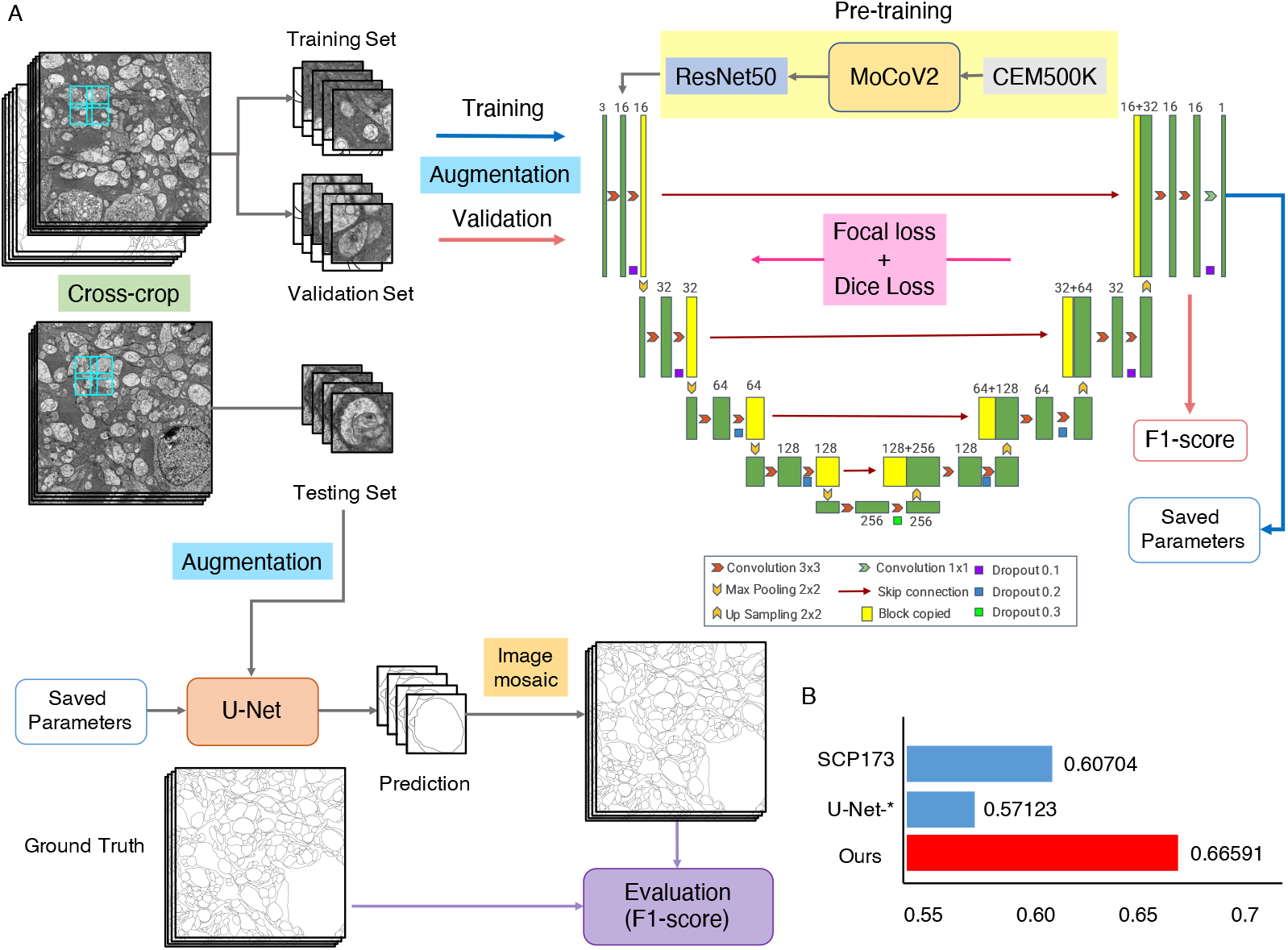
The U-Net-transfer method achieves the best performance on U-RISC. (A) The pretraining, training, and testing processing for U-Net. (B) The comparison of the F1-scores. “SCP-173” represents the top performance in the competition. U-Net-* represents the performance in ***Table 2***. Ours represents the performance of the U-Net-transfer method in this section.

## Discussion

This paper first proposed the U-RISC, a cell membrane EM dataset created through intensive and elaborate annotation. The dataset is characterized by the highest resolution and the largest single image size compared with other current publicly available annotated EM datasets. Next, we organized a segmentation competition on U-RISC and proposed the benchmark. During the competition, we noticed that the performances of popular deep learning methods were far below that of humans, which motivated us to explore the causes. Thus, we carried out a comprehensive survey on the deep learning methods participants applied in the competition. To our surprise, methods such as U-Net, LinkNet, and CASENet exhibited a significant drop of F1-score on U-RISC compared with ISBI 2012, from 0.9 to 0.6. To explore the mechanisms underlying this discrepant performance, we introduced a gradient-based attribution method, integrated gradient. Through attribution analysis of U-Net, we found the average pixel attribution field of U-RISC is larger than that of ISBI, corresponding to the size of cellular structure, and both of them are relatively small to the whole image size. By integrating currently available methods, we improve the benchmark to 0.67, about 10% higher than the top leader from the competition. Based on the analyses in this paper, here we raise some considerations about the challenges for deep learning-based segmentation algorithms brought by U-RISC and propose several suggestions for improving EM segmentation methods.

### Challenges for Deep Learning-based Segmentation

Benchmark showed that the segmentation performance of deep learning algorithms on U-RISC was still far behind the human level. U-RISC poses challenges for deep learning-based segmentation in the following aspects: (1) high computational costs needed to deal with large images, (2) the extreme sample imbalance caused by low ratio of cellular membrane pixels in the whole image, (3) side effects of typical data processing methods.

Deep learning itself is already a computationally intensive method. It would require more computational resources to process the images with a much larger size in U-RISC. In practical terms, taking U-Net as an example, processing a 1024 pixel × 1024 pixel image requires a GPU with 12GB memory. This memory is enough to deal with the images in ISBI 2012, of which the size is 512 pixel × 512 pixel. But the size of a single image in U-RISC is 9958 pixel × 9959 pixel, which is far beyond the processing ability of the commonly used 12 GB memory GPU. Therefore, the additional computational burden brought by U-RISC raises the first challenge for deep learning-based segmentation.

The problem of imbalanced samples widely exists in computational vision tasks (***Alejo et al., 2016; Li et al., 2010; Zhang et al., 2020***), which should be considered when designing algorithms. Cellular membrane segmentation is a typical situation of sample imbalance because cellular membrane only occupies a small proportion of the whole cell structure. According to statistics, the pixels belong to the cellular membrane account for 21.65% of the entire pixels of ISBI 2012. While the proportion in U-RISC is much smaller, 5.10%, making U-RISC an extremely imbalanced dataset. Pre-existing solutions were mainly proposed from several aspects: loss function design (***Lin et al., 2017; Cui et al., 2019***), data augmentation (***Yoo et al., 2020***), under/over-sampling (***Fernández et al., 2018; Yen and Lee, 2009***), and semantically multi-modal approaches (***Zhu et al., 2020***). However, even though the participants in the competition already used these approaches, the final results showed limited improvement of segmentation. So the imbalanced problem of U-RISC is yet to be solved and becomes another challenge for deep learning-based segmentation.

Proper data processing is essential and helpful to deep learning algorithms. For example, a downsampling process on raw images with an enormous size is commonly adopted in segmentation tasks (***Marin et al., 2019; Chen et al., 2014***). And in the Track 2 of our competition, we used the downsampled dataset to reduce computational consumption as usual. Surprisingly, we found that the F1-score of the same method dropped as well as the overall performance decreased in Track 2 compared with Track 1. We speculated a key reason might be the degradation of image quality from track 1 to Track 2. We confirmed the quality reduction through 4 representative indexes, including Brenner (***Subbarao and Tyan, 1998***), Variance (***Subbarao and Tyan, 1998***), SMD2 (***Thakkinstian et al., 2005***), and Vollath (***Vollath, 2008***) (shown in Appendix). More cautions should be paid when using traditional data processing methods, and more advanced data processing theories are expected from this point of view.

### Suggestions for the Improvement of Segmentation Methods

To some degree, increasing computational resources is a possible way to cope with the challenges mentioned above. However, it might not be easy for all the community researchers to access sufficient computational power; therefore innovations in algorithms are still crucial for our future success. To improve the performance of deep learning in EM segmentation, we provide several suggestions for developing deep learning algorithms from the following perspectives: model design, training techniques, data processing, loss function design, and visualization tools.

#### Model Design

As shown in the attribution analysis, the current models for segmentation, such as U-Net (***Ronneberger et al., 2015***), Efficientnet (***Tan and Le, 2019***), and CASENet (***Yu et al., 2017***), are designed to focus on local information to make predictions. However, in a high-resolution image, other structures, like organelle membrane and synaptic vesicles, might share similar features with the cellular membrane in a local scale, which leads to false-positive results. And this constitutes one of the major error types in the competition. Therefore, it might not be enough for the classifiers of a model to make correct decisions with only local features. And studies have shown that models using global information could improve performance greatly (***Liu et al., 2018, 2020; Wang et al., 2019***). Therefore, more global information could also be considered in the future design of the segmentation network.

#### Training Techniques

Skillful training techniques also can be helpful to improve the segmentation performance. According to our survey, a two-stage training strategy could be much better than a single-stage training strategy. A recent work also suggests that pretraining with domain-specific datasets can help network learning domain features (***Conrad and Narayan, 2021***). Besides that, much experience can be learnt from existing training methods. The Hypercolumns module (***Hariharan et al., 2015***) is used to accelerate the convergence of training by combining features at different scales, and the combination of features from different scales can help bring in global information. ScSE (***Roy et al., 2018***) module introduces attention mechanism into network, thus bringing in global information. Hybrid architectures can also be considered because of its ability to expand the receptive field (***Goceri, 2019***). In a word, improvement can be made at the phase of training by utilizing advanced training techniques.

#### Data Processing

Data processing is commonly used in deep learning, while traditional down-sampling methods were shown to have side effects in the competition. To alleviate the side effects, some quality enhancing methods for downsampled images could be expected, such as edge and region-based image interpolation algorithms (***Asuni and Giachetti, 2008; Hwang and Lee, 2004***), low bit rate-based approaches (***Wu et al., 2009; Lin and Dong, 2006***), and quality assessment researches (***Vu et al., 2018; Wang and Bovik, 2006; Wang et al., 2003***). Meanwhile, other data processing methods can also be taken into account. Take data augmentation as an example; by augmenting the training data randomly, the dependence of the model on specific attributes can be reduced, which can be beneficial in EM segmentation with many imbalanced samples.

#### Loss Function Design

Loss function design is another important part of deep learning. But many current loss functions have their own disadvantages in our competition. For example, Dice loss (***Dice, 1945***) was designed to optimize F1-score directly, without consideration of data imbalance. Focal loss (***Lin et al., 2017***) and BCE loss (***Cui et al., 2019***) were used in the competition to care more about data imbalance by giving different penalty according to sample difficulty, but the improvement was limited as the results have shown. A better design of loss function should take overall consideration of both sample imbalance and evaluation criteria. While the most common evaluation criterion, F1-score, a pixel-based statistic, is inconsistent with human subjective feeling to some extent. It might be a major cause of the performance gap between humans and algorithms. Some other structure-based criteria have appeared, such as V-Rand and V-info (***Arganda-Carreras et al., 2015***) integrating skeleton information of cell membrane, and ASSD (***Heimann et al., 2009***) considering the distance of point sets.

#### Visualization tools

Visualization tools can help us have a better understanding of the network. In this paper, with IG, we could learn the attribution fields of U-Net from the view of gradient, which inspires us to improve deep learning methods by paying more attention to global information. In comparison, many other visualization tools are starting from other characteristics of the network. Layer-wise relevance propagation (LRP) (***Bach et al., 2015***) and deep Taylor decomposition (DTD) (***Montavon et al., 2017***) get attribution distribution by modifying propagation rules. Information-based method IBA (***Schulz et al., 2020***) restricts the flow of information to accomplish attribution fields. Combining different visualization tools can help to promote much more insightful inspiration in improving deep learning methods.

Overall, we provide an annotated EM cellular membrane dataset, U-RISC, and its benchmark. It indeed brings many challenges for deep learning and promotes the development of deep learning methods for segmentation.

## Methods and Materials

### Dataset

The U-RISC dataset was annotated upon RC1, a large-scale retinal serial section transmission electron microscopic (ssTEM) dataset, publicly available upon request and detailedly described in the work of ***Anderson et al. (2011)***. RC1 came from the retina of a light-adapted female Dutch Belted rabbit after in vivo excitation mapping. The imaged volume represents the retinal tissue with a diameter of 0.25 mm, spanning the inner nuclear, inner plexiform, and ganglion cell layers. Serial EM sections were cut at 70-90 nm with a Leica UC6 ultramicrotome and captured at the resolution of 2.18 nm/pixel across both axes using SerialEM (***Mastronarde, 2005***). In RC1, there are in total 341 EM mosaics generated by the NCR Toolset (***Anderson et al., 2009***), and we clipped out 120 images in the size of 9958 pixel × 9959 pixel from randomly chosen sections.

To annotate cell membrane with high quality on 120 images, we launched an iterative annotation project that lasted for three months. All the annotators were trained to recognize and annotate cellular membrane in EM images, but only two-thirds of all, 53 annotators, were finally qualified to participate in the project according to their annotation results. In the iterative annotation procedure, each EM image would undergo three continuous rounds of annotation with the guidance of blind review. The final round of annotation was regarded as the “ground truth”. While since the first two rounds are valuable for analyzing the human learning process, we also reserved the intermediate results for public release. All of the U-RISC datasets are released at https://brain.baai.ac.cn/biodb-rabbit-details.html.

### Competition

The goal of the competition was to predict cell membranes in the EM images of U-RISC. Participants were required to return images depicting the boundary of all neurons. F1-score was selected as the evaluation criterion for the accuracy of the results. There are two tracks in the competition, images in Track 1 were kept as the original size (9958 pixel × 9959 pixel), images in Track 2 were downsampled to the size of 1024 pixel × 1024 pixel. Fifty images, 30 as the training dataset and 20 as the test dataset, were released in Track 1. And Track 2 contained 70 images in total, 40 training images, and 30 testing images. The training dataset included EM images with their corresponding ground truth, while the ground truth of the test dataset was kept private. In both tracks, ten images from the training dataset served as the validation dataset for the participants to monitor and develop their models. No statistical methods were used to determine the assignment of images in the whole arrangement.

### Segmentation Networks

We conducted experiments to compare the performance of the same methods on U-RISC (Track2) and ISBI 2012. Three representative deep learning networks were considered (***Table 2***), U-Net (***Ronneberger et al., 2015***), LinkNet (***Chaurasia and Culurciello, 2017***), and CASENet (***Yu et al., 2017***). The three networks are all pixel-based segmentation networks. To be specific, given the input image *x*, the goal of the networks is to classify the corresponding semantic cell membrane pixel by pixel. For the input image *x* and classification function *F*(*x*), *Y*{*p*|*X*,Θ}∈ [0,1] is taken as the output of the network, which represents the edge probability of the semantic category of the pixel *p*.Θ are the parameters in the network and are optimized in the training processes. Architectures of the three networks are described as follows.

### U-Net

U-Net (***Ronneberger et al., 2015***) is a classical fully convolutional network (that is, there is no fully connected operation in the network). The model is composed of two parts: contracting path and expansive path. The contracting path follows the typical architecture of a convolutional network. At each downsampling step, U-Net doubles the number of feature channels to gain a concatenation with the correspondingly cropped feature map from the contracting path. At the final layer a 1 ×1 convolution is used to map each 64-component feature vector to the desired number of classes. In total the network has 23 convolutional layers. We use ResNet50 as its encoder.

### LinkNet

The model structure of LinkNet (***Chaurasia and Culurciello, 2017***) is almost similar to U-Net, which is a typical encoder-decoder structure. The encoder starts with an initial block which performs convolution on input image with a kernel of size 7×7 and a stride of 2. This block also performs spatial max-pooling in an area of 3×3 with a stride of 2. The later portion of encoder consists of residual blocks and is represented as the encoder-block. To reduce parameters, LinkNet uses ResNet18 as its encoder.

### CASENet

CASENet (***Yu et al., 2017***) is an end-to-end deep semantic edge learning architecture adopting ResNet-101 as backbone. The classification module here consists of a 1×1 convolution and a bilinear interpolation up-sampling layer to generate *M* active images, each image size is the same as the original image. Each residual block is followed by a classification module to obtain five classification activation graphs. Then, a sliced concatenation layer is used to fuse the *M* classification activation graphs, and finally a 5 × *M* channel activation graph is obtained. The activation graphs are used as the input of the fused classification layer to obtain a *M*-channel activation graph. The fusion classification layer is convolution of *M* group 1×1.

### Transfer learning

The pre-trained model from (***Conrad and Narayan, 2021***) was used in our method, specifically MoCoV2 (***He et al., 2020***) and CEM500K (***Conrad and Narayan, 2021***) were respectively selected as the pretraining method and dataset.

### Training settings

For each dataset, same training and testing data distribution was utilized on the three methods. For U-RISC, during the training, the original images were cut into 1024 × 1024 patches with overlaps. And the patches were randomly assigned into the training set and balidation set according to the ratio of 50,000/20,000. For ISBI 2012, 20 images were used for training and 10 images were used for testing.

### Loss function and optimization

For each algorithm, we used the same loss function and optimization method. Specifically, focal loss and dice loss were chosen. Define *y* as the ground truth segmentation and 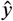 as the predicted segmentation. The calculations of focal loss and dice loss were:

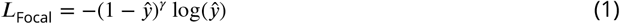

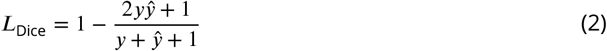

And the final loss function is the summation of the two losses with the proportion of 1: *λ*. That is *L* = *L*_Focal_+*λL*_Dice_. We set *λ* = 1 in these experiments. When optimizing the parameters in the network, we chose Adam (***Kingma and Ba, 2014***) as the optimizer.

### Parameter settings

Data augmentation (random horizontal/vertical flip, random rotation, random zoom, random cropping, random cropping, random translation, random contrast, and random color jitter) were used. Four Nvidia V100 GPUs were used for training. In the testing stage, the original images were cut into the same size as the training images, and the patchs were tested. These patches were eventually mosaiced back to the original size for evaluation. The parameters settings are shown in ***Table 3***. Mean value and standard error are computed over testing images of each dataset. The methods with “_*” in the table represent that they are implemented by ourselves.

**Table 3.**
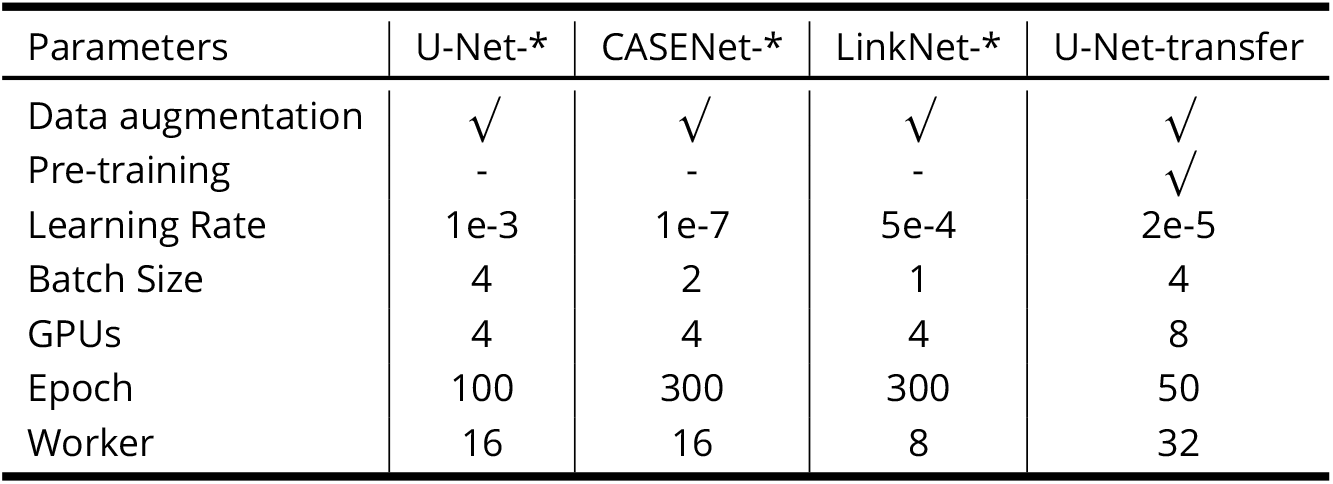
Parameter settings.

### Image definition criteria

Four representative image definition criteria, **Brenner**(***Subbarao and Tyan, 1998***),**SMD2**(***Thakkins-tian et al., 2005***),**Variance**(***Saltelli et al., 2010***), and **Vollath**(***Vollath, 2008***) were used for analyzing the effects of downsampling on EM images. The former two consider the difference and variance of gray values between adjacent pixels, while the latter two consider the whole image.

Brenner gradient function simply calculates the square of the gray difference between two adjacent pixels.

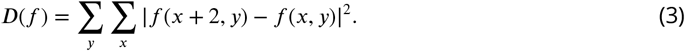

where: *f* (*x, y*) represents the gray value of pixel (*x, y*) corresponding to image *f*, and *D*(*f*) is the result of image definition calculation (the same below).

SMD2 multiplies two gray variances in each pixel field and then accumulates them one by one.

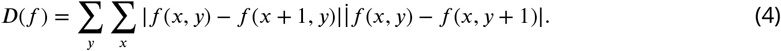

The Variance function is defined as

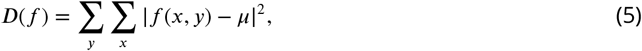

where: *μ* is the average gray value of the whole image, which is sensitive to noise. The purer the image, the smaller the function value.

The Vollath function is defined as

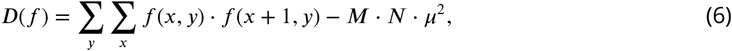

where: *μ* is the average gray value of the whole image, *M* and *N* are the width and height of the image respectively.

### Attribution analysis

The purpose of the integral gradient (IG) method is to quantify the contribution of each part of the input feature to the decision. For a given input image *x* and model *F*(*x*), the goal of the network is to find out which pixels or features in *x* have an important influence on the decision-making of the model or sort the importance of each pixel or feature in *x*. Such a process is defined as attribution. IG uses the integral value along the whole gradient line from input to output. In the cell membrane segmentation task, for the decision of a pixel of *y* (predicted as the cell membrane or not), we can get the contribution of each pixel of the input image. Put the contribution of each pixel together, we record it as an ***Attribution Field** A*, whose size is the same as the original image. And the value of each pixel is *w_i,j_*, representing the contribution decision of pixel *x_i,j_* to *y*. *w_i,j_* is normalized to [−1,1].

In binary segmentation task, for the current input image *x*, if we know that the output *y* is a specific value, such as *y* = 0, and the corresponding reference image is 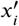, then we can take a linear interpolation, that is

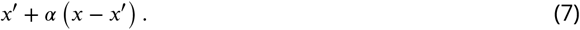

If the constant *α* = 0, then the input image is the base image as 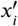. And if *α* = 1, then the input image is the current image, which is *x*. When 0 ≤ *α* ≤ 1, it can be other images.

For the output of the neural network *F*(*x*), the formula of the IG method is shown as

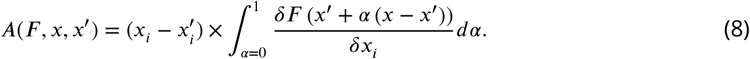

In formula 8, *δx_i_* on the denominator denotes variation. This design makes the whole partial derivative transform into the form of variation. The variation boundary is the reference image and the current image. The integral gradient method uses linear interpolation as the variational path.

That is,

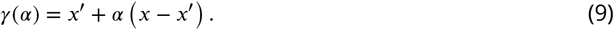

Here we select the random noise image as the reference image.

As the resolution and image size of U-RISC and ISBI 2012 are different, for a fair comparison, we define the size of **Pixel Attribution Field** as *S_k_*, which represents the physical size corresponding to the pixel area with fixed contribution value threshold *k*. If the contribution value *w_i,j_* is greater than *k*, the pixel is the one with higher contribution to decision-making. Area of attribution field *S_k_* is obtained by multiplying the areas (attribution value *w_i,j_* > *k*) and the conrresponding physical size of a pixel (square of resolution *h*).

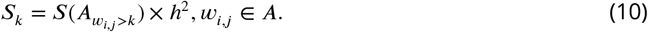

### Data analysis

All statistical tests used, including statistic values and sample sizes, are provided in the figure captions, including mean and standard. All analyses were performed using custom software developed using the following tools and software: MATLAB (R2018a), Python (3.6), PyTorch (1.6.0), NumPy (1.19.0), SciPy (1.5.1), and matplotlib (2.2.3).

## Acknowledgments

We thank Bryan W. Jones for kindly providing RC1 connectomic dataset. This project is supported by Cognitive Computing Grant No.62088102.

## Appendix 1 The Effects of Downsampling on U-RISC Dataset

To our best knowledge, downsampling is an effective approach to process images with enormous size in segmentation tasks (***Marin et al., 2019; Chen et al., 2014***). With the purpose of investigating the effects of downsampling on image definition quality, we analyzed the gray-scale histograms on a group of original and downsampled images in U-RISC and then calculated the definition values of images in Track 2.

We first found that downsampling produced numerous sharp changes between adjacent gray values (***Figure 1***(C)). In our analysis, the mutations might engender the texture information in the image to be blurred. For example, the bilayer structure of the membrane disappeared after downsampled (***Figure 1***(A,B)), and some of the cell membranes which were hard to recognized became obscured. We then found that the defining quality of the images in Track 2 became lower after downsampling. Four definition indexes of all the images in Track 2 were calculated. Brenner (***Subbarao and Tyan, 1998***), Variance (***Subbarao and Tyan, 1998***), SMD2 (***Thakkinstian et al., 2005***), and Vollath (***Vollath, 2008***) are common indexes to show the gray value change between adjacent pixels. Results suggested that image indexes were significantly decreased after downsampling (***Figure 1***(D)), and all the four indexes dropped about ten times. Therefore, the analysis indicated that images were heavily blurred after the downsampling operation.

**Appendix 1 Figure 1.**
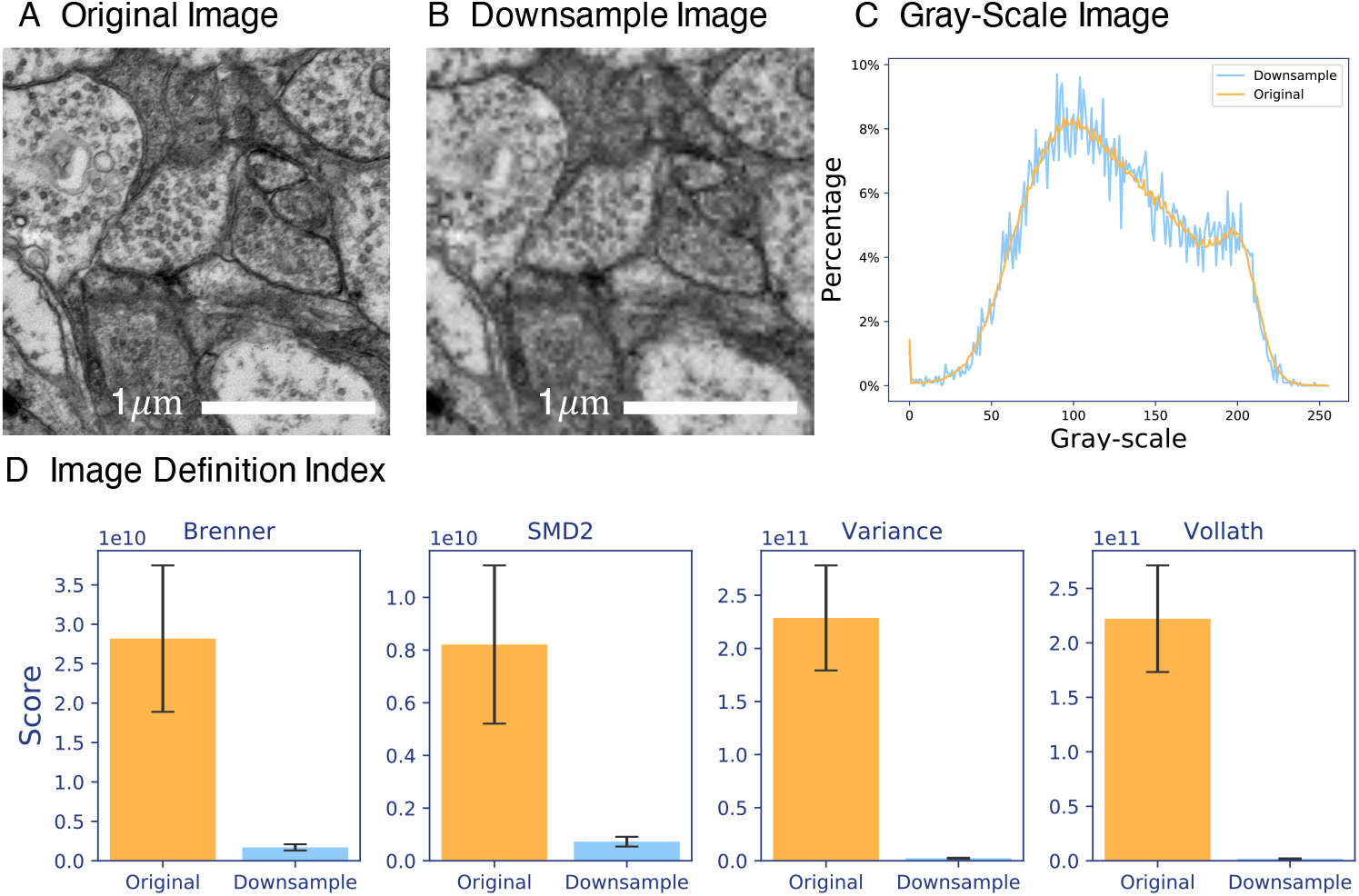
Differences between original image and downsampled image.(A) The crop of original image. (B) The crop of downsampled image at the same position. The Gray-scale Histograms is calculated on A and B. (C) The scores of definition indexes calculated on the whole U-RISC dataset before and after downsampling. Details of indexes are described in Methods and Materials.

---

**Figure 6-Figure supplement 1.**
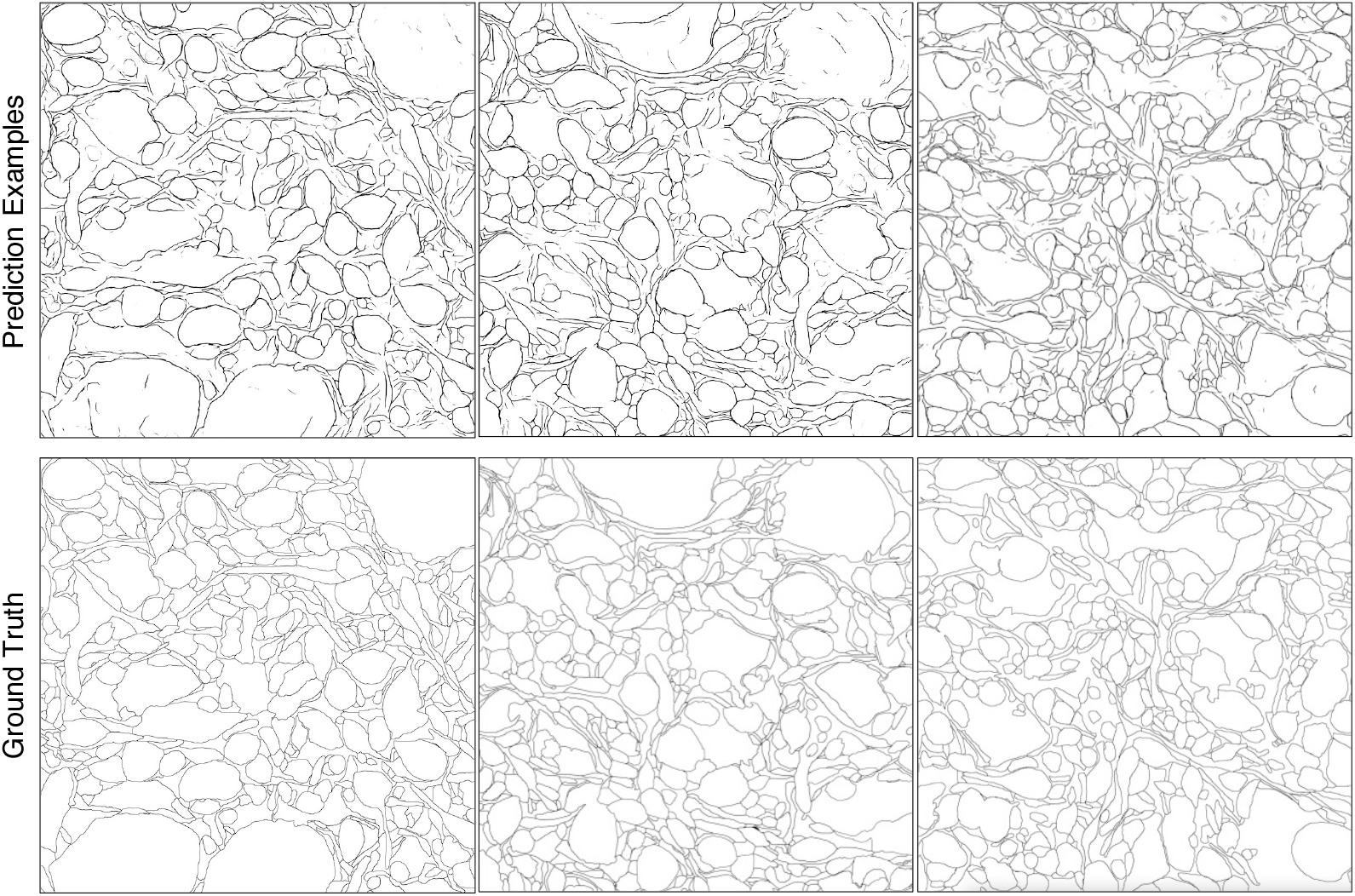
Supplements for segmentation predictions of U-RISC.

**Figure 6-Figure supplement 2.**
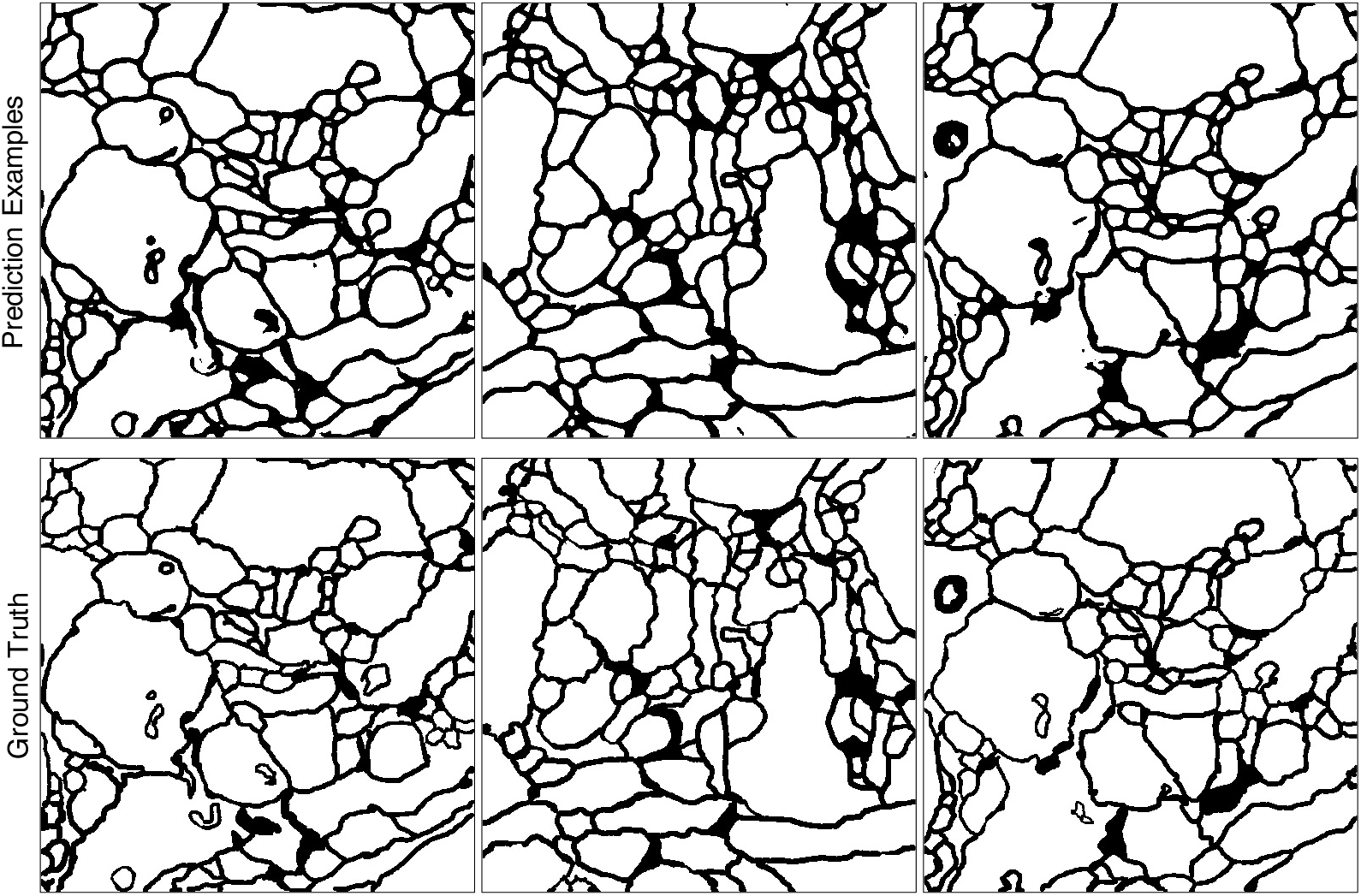
Supplements for segmentation predictions of ISBI 2012.

**Figure 8-Figure supplement 1.**
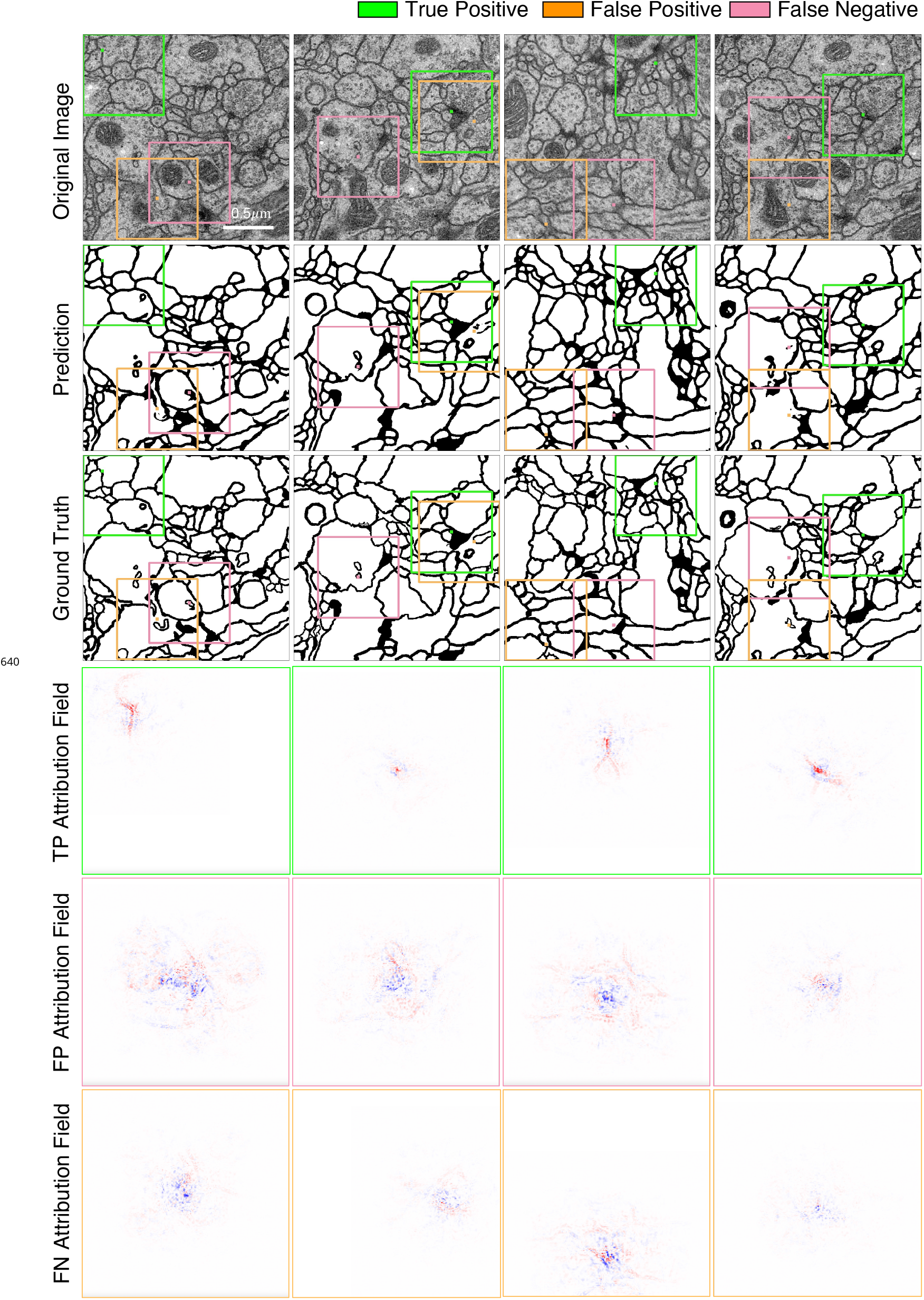
Supplements for attribution analysis on ISBI 2012.

**Figure 8-Figure supplement 2.**
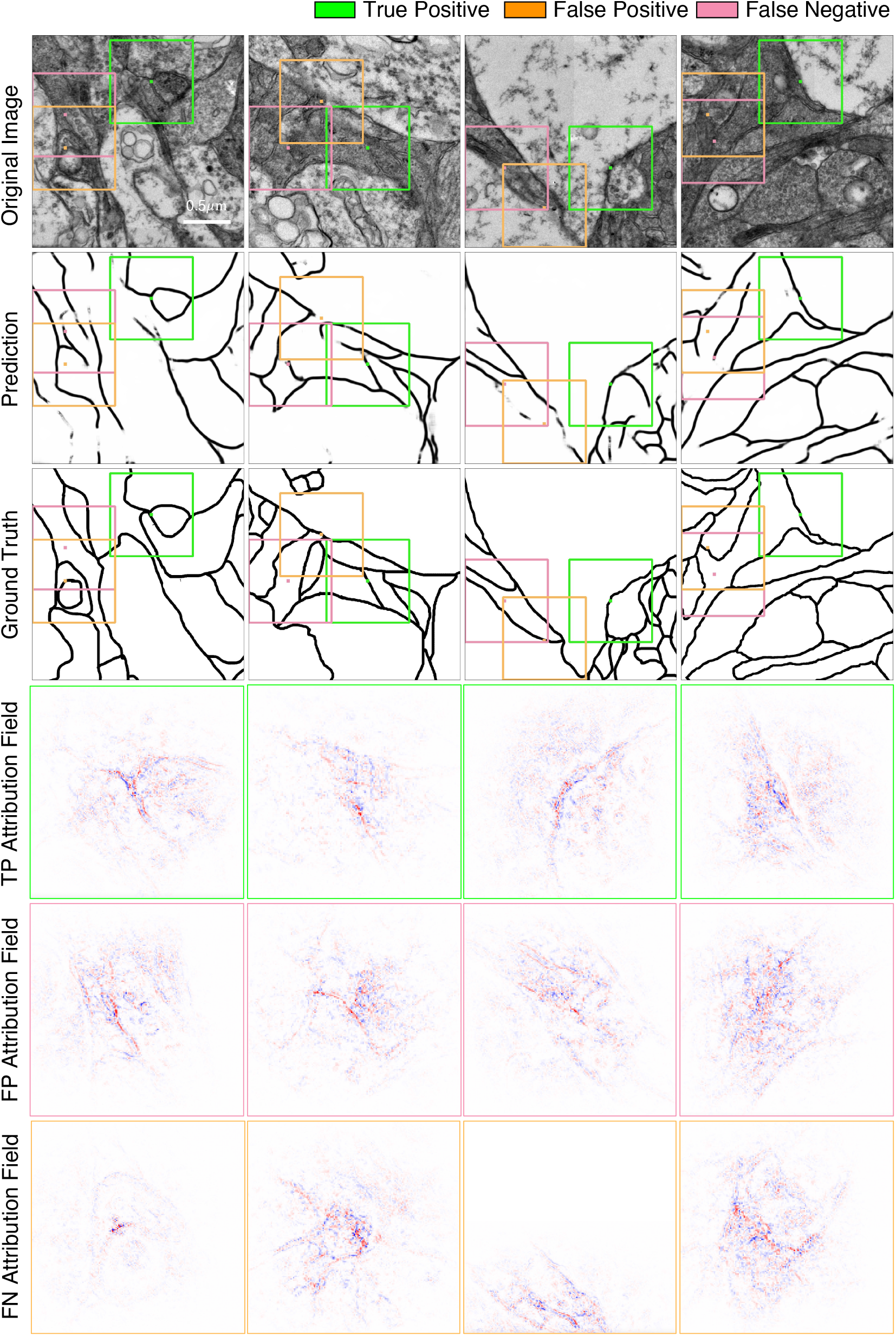
Supplements for attribution analysis on U-RISC.

## Footnotes

1 https://www.biendata.xyz/competition/urisc/

1 Tencent Holdings Ltd (China)

2 University of Chinese Academy of Sciences (China)

3 Sichuan University (China)

4 Nanjing University (China)

5 Hangzhou Dianzi University (China)

6 University of Science and Technology of China

7 Tsinghua University (China)

8 Jilin University (China)

